# *In vivo* activation of the *dia* BGC allows consolidation of the biosynthetic pathways of diaporthin, dichlorodiaporthin, diaporthinic acid, and diaporthinol

**DOI:** 10.1101/2025.03.31.646288

**Authors:** Isabella Burger, Simon Leonhartsberger, Kathrin Peikert, Lukas Fourtis, Polina Atanasova, Lara T.S. Kramer, Richard Fried, Christian Stanetty, Florian Rudroff, Ruth Birner-Gruenberger, Robert L. Mach, Astrid R. Mach-Aigner, Matthias Schittmayer, Christian Zimmermann

**Author notes:** IB and SL contributed equally to this study.

## Abstract

Fungal secondary metabolites exhibit remarkable chemical diversity and biological activity, making them valuable sources of bioactive compounds. A group of polyketides, with a similar core structure, have been isolated previously from different fungi - the phytotoxins diaporthin and orthosporin, the mediocre antimicrobial dichlorodiaporthin, as well as diaporthinol and diaporthinic acid. Previous studies in *Aspergillus oryzae* suggest that diaporthin and orthosporin originate from a different biosynthetic gene cluster than dichlorodiaporthin, while the biosynthetic routes for diaporthinic acid and diaporthinol have not yet been identified.

In this study, we successfully activated the *dia* biosynthetic gene cluster in *Trichoderma reesei* via transcription factor overexpression, leading to the identification of diaporthinic acid as the primary metabolite. Structural elucidation using NMR confirmed its identity, while bioactivity assays revealed no significant antimicrobial effects. Further, diaporthin, orthosporin, dichlorodiaporthin, and diaporthinol, as well as an isomer of alternariol were attributed to the same cluster. Gene deletion experiments demonstrated that Dia1, Dia4, and Dia5 are essential for diaporthinic acid biosynthesis, with Dia4 likely catalyzing the oxidation of dichlorodiaporthin yielding diaporthinic acid. Dia2 and Dia3 were found to be dispensable, challenging previous *in vitro* findings for the biosynthesis of dichlorodiaporthin. These findings provide new insights into the biosynthetic network of the *dia* BGC, expanding our understanding of natural product biosynthesis in filamentous fungi.

## Introduction

Fungal secondary metabolites are an important source of pharmaceuticals and other useful substances. Based on their vast chemical diversity, they possess a large number of bioactive properties ^1,2^. The best-known example is probably antibiotics, with penicillin as a society-changing discovery ^3^. Humankind has proceeded to study the fungal secondary metabolism in the hope of finding new substances and also to understand their biosynthetic pathways and the properties of the enzymes involved ^4,5^.

The enzyme AoiQ sparked scientific interest due to its unusual nature. It is a bifunctional enzyme containing a halogenase domain and a methyltransferase domain^6^. Chankhamjon *et al*. first described that this enzyme is essential for the formation of dichlorodiaporthin (**1**) in *Aspergillus oryzae*. The encoding gene is found in a biosynthetic gene cluster (BGC), which also contains the genes for a non-reducing polyketide synthase (PKS), a β-lactamase-like enzyme, a short-chain dehydrogenase/reductase (SDR), and a flavin-dependent monooxygenase (FMO)^6^. AoiQ and the *dia* BGC are found in several fungi (Fig. 1A and Table 1), including *Aspergillus oryzae, A. nidulans, Trichoderma harzianum*, and *T. virens* and *T. reesei*. The BGC of *Penicillium nalgiovense* contains two distinct genes encoding for separate halogenase and methyltransferase (Fig. 1A)^7^.

**Table 1.** Genes in the *dia* BGC (see also Additional File1 for the corresponding genbank files)

| Gene name | Protein ID | Enzyme class | Homolog in *A. oryzae* | Homolog in *A. nidulans* | Homolog in *P. nalgiovense* |
| --- | --- | --- | --- | --- | --- |
| *dia1* | 81964 | polyketide synthase | *diaA* | *pkgA* | *ngvA* |
| *dia2* | 69625 | beta-lactamase-like | *diaB* | *pkgB* | *ngvB* |
| *dia3* | 123964 | dehydrogenase | *diaC* | *pkgC* | *ngvC* |
| *dia5* | 69652 | bifunctional flavin-dependent halogenase / methyltransferase | *aoiq* | *aoiq* | *ngvE* (FDH)<br>*ngvF* (MT) |
| *dia4* | 69650 | FAD-dependent oxidoreductase | *diaD* | *pkgD* | *ngvD* |
| *diaR1* | 111742 | zinc cluster protein | XM_003190268.1 | ANIA_07073 | n/a |

**Figure 1.**
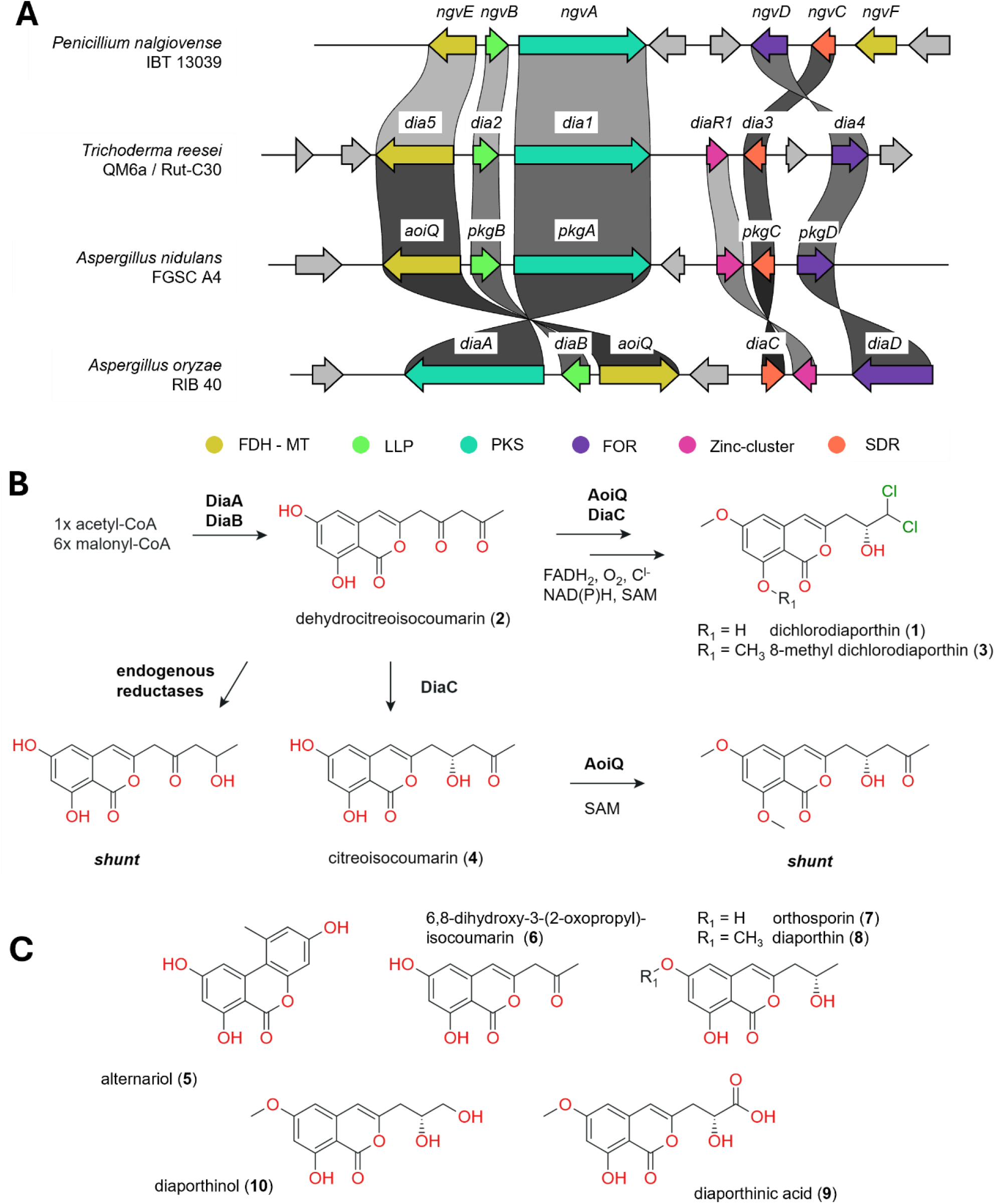
(**A**) The *dia* BGC and its homologs from the indicated fungal strain were compared and visualized using the cblaster and the clinker tool^10^. FDH-MT, flavin-dependent halogenase-methyltransferase; LLP, beta-lactamase-like protein; PKS, polyketide synthase; FOR, FAD-dependent oxidoreductase; Zinc-cluster (protein); SDR, short chain dehydrogenase/reductase. (**B**) The previously described biosynthetic pathway towards 8-methyl dichlorodiaporthin according to Liu *et al*.7. (**C**) Other diaporthin-related, reported metabolites.

In 2022, Liu *et al*. studied the enzyme AoiQ in detail and scrutinized its role and precise mode of action during the biosynthesis of dichlorodiaporthin (**1**)^7^. They observed that AoiQ is able to catalyze gem-dichlorination of 1,3-diketones. To gain insight into the biosynthetic pathway of the *dia* BGC, Liu *et al*. heterologously expressed *the A. oryzae* enzymes DiaA-C and AoiQ in *Saccharomyces cerevisiae* and performed several *in vitro* assays with AioQ and DiaC produced in *Escherichia coli*. Based on their results, Liu *et al*. proposed the following biosynthetic pathway (Fig. 1B). DiaA and DiaB are responsible for the formation of dehydrocitreoisocoumarin (**2**). As the PKS does not contain an enoyl-reductase domain, which is normally involved in release of the polyketide, the β-lactamase-like enzyme is suggested to take over this role. DiaC and AoiQ then act simultaneously resulting in the formation of 8-methyl dichlorodiaporthin (**3**). They also reported on several shunt products, including citreoisocoumarin (**4**). When AoiQ was substituted with the homologs from *P. nalgiovense, A. nidulans*, or *T. harzianum* in the yeast assays, dichlorodiaporthin (**1**) was produced, suggesting that only the methyltransferase domain of *A. oryzae* AoiQ is able to catalyze the *O*-8 methylation^7^.

In another previous study, the DiaA and DiaB homologs of *A. nidulans* (PkgA, ANID_07071.1 and PkgB, ANID_07070.1) were overexpressed in the native host^8^. Ahuja *et al*. reported on the detection of compounds with masses corresponding to citreoisocoumarin (**4**) and alternariol (**5**) (Fig. 1C)^8^. In contrast, Liu *et al*., did not report on the formation of **5**, but found small amounts of **4**, which was attributed to the activity of DiaC on **2**^7^. Importantly, the flavin-dependent monooxygenase DiaD or its homologs were not yet included in any study, and its activity remains unknown. Further, there are no reports on a fully activated *dia* BGC *in vivo* in any fungus to date.

Notably, *A. oryzae* contains a similar BGC, the *aoi* BGC, which contains a PKS, an oxygenase, and a methyltransferase ^9^. This BGC was activated previously by a transcription factor over expression strategy^9^. Nakazawa *et al*. suggested that the PKS AoiG synthesized 6,8-dihydroxy-3-(2-oxopropyl)-isocoumarin (**6**), which was reduced by the oxidoreductase AoiI yielding orthosporin (**7**). **7** was suggested to be further methylated by AoiF resulting in diaporthin (**8**) based on elevated transcript levels for these enzymes in the activated strain (Fig. 1C). Later, Chankhamjon et al. suggest that the methylation of **7** might be catalyzed by AoiQ based on *in vitro* assays^6^. However, **1** and **8** are considered to be the products of two distinct BGCs in *A. oryzae*^6,7^.

In this study, we activated the *dia* BGC in *T. reesei* using a transcription factor overexpression strategy as previously used to activate the ilicicolin H BGC^11^. We isolated and identified the main product of the *dia* BGC, diaporthinic acid (**9**) via NMR, and tested it for antibacterial and antifungal properties. We further deleted all *dia* genes individually in the activated strain and performed HPLC-MS/MS analyses to scrutinize the *in vivo* biosynthetic pathway. Additionally, we measured the transcript levels of the genes in all strains. Finally, the gene *dia3* was also deleted in the wild-type background to test its influence on physiology of *T. reesei* within and outside of the *dia* biosynthetic pathway.

## Results

### Overexpression of DiaR1 leads to activation of the *dia* BGC

The *dia* BGC in *T. reesei* contains a gene encoding for a zinc cluster protein (protein ID 111742^12^, see Fig. 1 and Table 1). Notably, the predicted protein consists of only 309 amino acids and the NCBI conserved domain search^13,14^ only identifies a Gal4-like DNA binding domain (smart00066) at the N-terminus (R12 – R52) but no transactivation domain. This is in stark contrast to the standard model for Gal4-like TFs, which entails the presence of both domains in a single protein^15^. Regardless, we decided to overexpress the zinc cluster protein in *T. reesei* by putting the coding region under the control of the constitutive *tef1* promoter and inserting it in front of the *pyr4* gene as described in Derntl and Kiesenhofer et al.^16^. In the resulting strain, *T. reesei* OEdiaR1, we could detect elevated transcript levels of all genes in the *dia* BGC in comparison to the control strain QM6a Δmus53 (“WT”) (Fig. 2, Additional File 2). The transcript levels for *dia3* are only moderately upregulated, because this gene is already transcribed in the control strain

**Figure 2.**
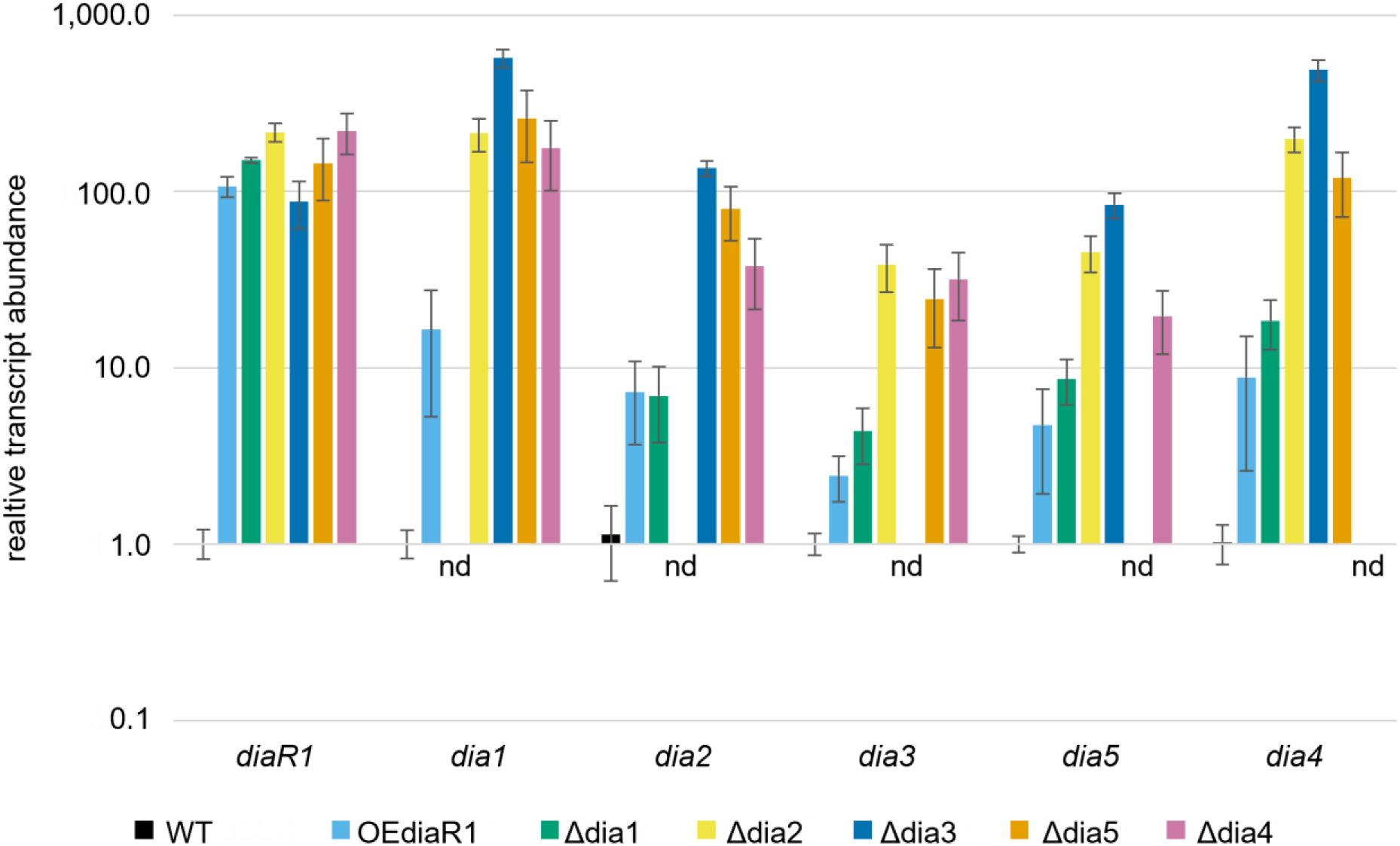
The indicated strains were cultivated in defined medium and the total RNA extracted after 48 hours. The relative transcript abundances of the indicated genes were measured in the indicated strains via a RT-qPCR analysis and normalized to the transcript levels in the WT strain. nd, not detected. The values are the means of biological quadruplicates. The error bars represent standard deviation.

(Ct approx. 23). Other genes with appreciable transcript levels in the control strain are *dia5* and *diaR1* (Ct approx. 24.5 and 26.5, respectively), whereas we could not detect transcript levels distinguishable from the negative control for the rest of the *dia* genes (Additional File 2).

### The main compound of the *dia* BGC is diaporthinic acid

Upon successful activation of the *dia* BGC, we performed a UV-vis spectrometric screen of the supernatant and compared it to the control strain. We observed several absorption peaks in the spectrum of the OEdiaR1 strain with local maxima at 238 nm, 244 nm, 279nm, and 330nm (Fig. S1A), indicating the presence of at least one compound that is not present in the control strain. We observed several peaks in the chromatogram during a HPLC-PDA/MS analysis of the supernatant (Fig. S1B). Purification of the crude mixture via preparative RP-HPLC allowed the isolation of the substance corresponding to the indicated peak with *m/z* (ESI+) = 281 and *m/z* (ESI-) = 279. We could identify the compound as diaporthinic acid (**9**) in subsequent 1D- and 2D-NMR experiments (Fig. S2).

Next, we performed a minimal inhibitory concentration (MIC) assay to test if **9** possesses antimicrobial properties. In our experimental setup, we could not detect any growth inhibitory effects against *E. coli, Bacillus subtilis, S. cerevisiae*, or *A. nidulans* in concentrations up to 128 µg/ml (Fig. S3, Additional File 3).

### Dia1, Dia5, and Dia4 are essential for diaporthinic acid production

As mentioned, the *dia* BGC was linked to citreoisocoumarin (**4**), dicholorodiaporthin (**1**) and 8-methyl-dichlorodiaporthin (**3**) in previous studies. Diaporthinic acid (**9**), on the other hand, was only isolated from different fungi, e.g. *P. nalgiovense* (strains IBT 12679, IBT 13296, and IBT 13330)^17^ or *Didymella pomorum* (previously *Phoma pomorum*)^18^ but has not yet been assigned to a BGC. To test whether **9** is in fact originating from the *dia* BGC, we first deleted the *dia1* gene in the OEdiaR1 strain. This gene encodes for a PKS-NRPS hybrid enzyme which is the core enzyme of the BGC according to the literature^7^. The resulting strain still had an active BGC, with transcript levels of all genes similar to the OEdiaR1 strain (Fig. 2, Additional File 2), but we could not detect any of the absorption peaks in the UV-vis spectrum (Fig. S1A), nor **9** when conducting a HPLC-MS/MS analysis (Fig. 3). This strongly indicates that the activity of the PKS-NRPS Dia1 is essential for the biosynthesis of **9** and thus that **9** originates from the *dia* BGC.

**Figure 3.**
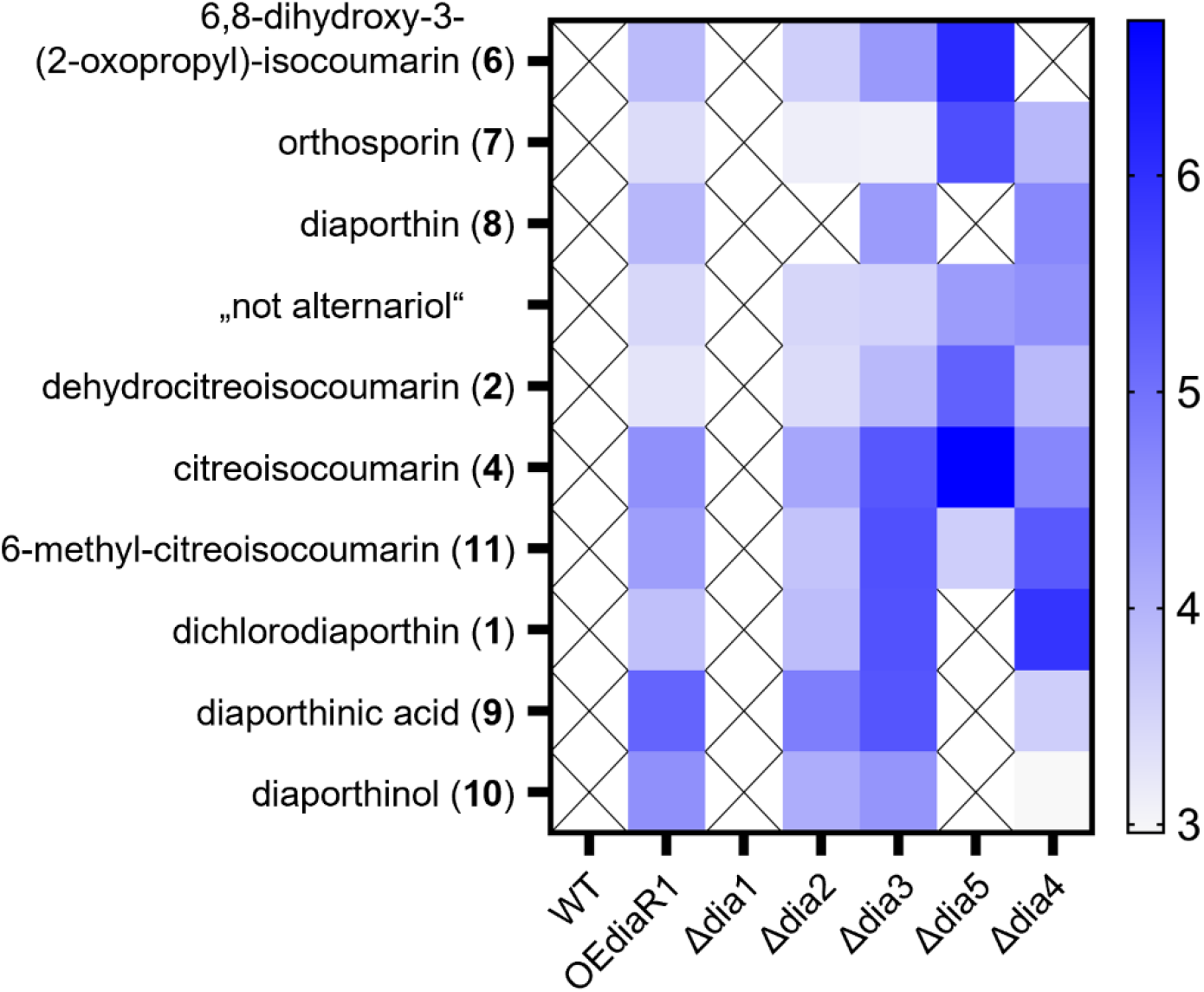
The indicated strains were cultivated in defined medium for 72 hours, followed by untargeted metabolomics analysis of their culture supernatants. The heatmap depicts the TIC-normalized and log_10_ transformed peak areas of the detected compounds.

To gain further insight into the biosynthetic pathway of **9**, we deleted all of the remaining genes of the *dia* BGC individually in the OEdiaR1 strain. Like in the *dia1* deletion strain, we could still detect high transcript levels of all *dia* genes except the respective deleted genes. Notably, the transcript levels were higher in these strains than in the OEdiaR1 and the *dia1* deletion strain; with the highest levels in the *dia3* deletion strain (Fig. 2, Additional File 2).

These generated strains were subjected to untargeted metabolomics measurements by HPLC-MS/MS. Notably, NMR verified standards were only available for **5** and **9**. Identification of the other compounds was based on exact mass only, which has to be considered when interpreting the results. Please refer to Table 2 for a list of all detected compounds used for data interpretation in this study.

**Table 2.** Detected analytes, including sum formula and exact mass information.

| Assumed compound | Sum formular | Exact mass [Da] | Matching to standard? |
| --- | --- | --- | --- |
| dichlorodiaporthin (**1**) | C<sub>13</sub>H<sub>12</sub>Cl<sub>2</sub>O<sub>5</sub> | 318.0062 | n/a |
| dehydrocitreisocoumarin (**2**) | C<sub>14</sub>H<sub>12</sub>O<sub>6</sub> | 276.0634 | n/a |
| citreisocoumarin (**4**) | C<sub>14</sub>H<sub>14</sub>O<sub>6</sub> | 278.0790 | n/a |
| alternariol (**5**) | C<sub>14</sub>H<sub>10</sub>O<sub>5</sub> | 258.0528 | NO |
| 6,8-dehydroxy-3-(2-oxopropyl)-isocoumarin (**6**) | C<sub>12</sub>H<sub>10</sub>O<sub>5</sub> | 234.0528 | n/a |
| orthosporin (**7**) | C<sub>12</sub>H<sub>12</sub>O<sub>5</sub> | 236.0685 | n/a |
| diaporthin (**8**) | C<sub>13</sub>H<sub>14</sub>O<sub>5</sub> | 250.0841 | n/a |
| diaporthinic acid (**9**) | C<sub>13</sub>H<sub>12</sub>O<sub>7</sub> | 280.0583 | YES |
| diaporthinol (**10**) | C<sub>13</sub>H<sub>14</sub>O<sub>6</sub> | 266.0790 | n/a |
| 6-methyl-citreisocoumarin (**11**) | C<sub>15</sub>H<sub>16</sub>O<sub>6</sub> | 292.0947 | n/a |

In accordance with previous results, the deletion of *dia5* (homolog of *aoiQ*) resulted in an abolishment of compound **1** production (Fig. 3). In this strain, we could not detect signals for **8, 9**, or **10**, but observed an accumulation of compounds with masses matching **2, 4, 6**, and **7** compared to the OEdiaR1 strain (Fig. 3). The deletion of *dia4* drastically reduced the abundance of **9** and the compound with a matching mass for **10** and led to a strong accumulation of the assumed **1**, and to a lesser degree also the assumed **11** (Fig. 3), suggesting that **1** might be the substrate for Dia4.

In both strains, Δdia4 and Δdia5, we observed a slightly stronger signal for a compound matching the mass of **5**. However, despite exhibiting a matching exact mass (*m/z* = 259.0614, theoretical [M+H]+ 259.0601), comparison with the commercially available standard revealed a significant retention time and fragmentation pattern mismatch. Hence, this compound is not **5**, and we designated it as “not alternariol”. Nonetheless, the compound can still be attributed to the *dia* BGC, as it was not detected in the Δdia1 strain (Fig. 3).

### Dia2 and Dia3, essential for dichlorodiaporthin formation *in vitro*, are dispensable *in vivo*

Previously, Liu *et al*. demonstrated that *A. oryzae* DiaA and DiaB work together to synthesize **2** and that DiaC and AioQ act simultaneously on **2** yielding **3**^7^. As mentioned, **3** differs from **1** by a methyl group that is uniquely added by the *A. oryzae* AoiQ but none of its homologs from other fungi^7^; **3** from *A. oryzae* thus equals **1** in the *T. reesei* biosynthetic pathway. Our results suggest that neither Dia2 nor Dia3 are essential for the biosynthesis of **1** *in vivo*. The deletion of *dia2*, lowered the signal intensities for the compounds with masses matching nearly all anticipated metabolites in comparison to the OEdiaR1 strain (Fig. 3). Only the signal for compound **8** was not detected anymore (Fig. 3). The deletion of *dia3* did not result in an abolishment of **1** or **9** production in *T. reesei* (Fig. 3), or any other intermediate. We even observed substantially stronger signals for the assumed **1** and **11** in the Δdia3 strain compared to the OEdiaR1 strain.

At this point, we want to mention that the accumulated biomass of the Δdia3 strain is strongly reduced in comparison to the other strains (Fig. S4). As mentioned above, *dia3* is also expressed in the wild-type strain (Fig. 2, Additional File 2), indicating that Dia3 is not exclusively participating in the biosynthetic pathway of the *dia* BGC, but might also catalyze other reactions. Thus, we sought to determine whether the reduced growth of Δdia3 is specifically due to the absence of Dia3 in the biosynthetic *dia* pathway or a consequence of its overall loss. Therefore, we deleted the *dia3* gene in the wild-type strain. The resulting strain WT/Δdia3 has no growth defects and grows similarly to the wild-type strain (Fig. S4). This strongly indicates that Dia3 plays an important role during the biosynthesis of **1** and **9**, but this is not necessarily its previously reported catalytic activity.

## Discussion

Previously, **9** was first isolated together with **1** and **10** from three isolates of *P. nalgiovense* (IBT 12679, IBT 13296, and IBT 13330) as described by Larsen *et al*.^17^. Also *D. pomorum* was described as a producer of **9**. Sørensen *et al*. detected **9** together with **1, 4, 8, 10**, and **11** in 27 of 27 tested strains from Denmark, the Netherlands or Gemany^18^. Importantly, these studies only described the presence of **9** but did not link **9** to a BGC. Using a BGC activation strategy, we could attribute **9** to the *dia* BGC in this study. The genomes of the *P. nalgiovense* and *D. pomorum* strains used in the previous studies are not available, but we found homologs of the *dia* BGCs in the genomes of sequenced strains using the CompArative GEne Cluster Analysis Toolbox^10^. Exemplarily, the BGC of the *P. nalgiovense* strain IBT 13039 is included in Fig. 1. Homologs of *T. reesei* Dia1-5 are found in the genome of the Canadian isolate *D. pomorum* M27-16 on scaffold 121 (GenBank: JAKJXN010000121.1) (Table S1). Taken together, we postulate that the *dia* BGC is responsible for the production of **9** not only in *T. reesei* but also in *P. nalgionvense, D. pomorum*, and other fungi that contain this BGC^7^ (Fig. 1, Table 1; Table S1).

In this study, we observed that the main product of the *dia* pathway is **9**, which accumulates in extraordinary abundance in the OEdiaR1 strain in relation to the other metabolites. The deletion of *dia5* (homolog of *aoiQ*) resulted in a complete abolishment of **9** and also **3, 8**, and **10**. The absence of **3** and **8** is in accordance with previous results. Liu *et al*. suggest that AoiQ is essential for the production of **8** and **1** in *A. oryzae* (**1** corresponds to **3** in *T. reesei*)^7^. The absence of **9** and **10** in this strain also puts them downstream of Dia5. In the Δdia4 strain, we detected substantially lower amounts of **9** compared to OEdiaR1 and a strong accumulation of **1**, which suggests that **1** is the substrate for Dia4. Dia4 contains a p-cresol methylhydroxylase (PCMH)-type FAD-binding domain (Uniprot ID G0RVA9). This domain is found in various oxidases and dehydrogenases according to its Interpro entry (IPR016166). Dia4 is therefore a suitable candidate for the oxidation of **1** yielding **9** in *T. reesei*. The proposed biosynthetic pathways based on our results and previous results from the *A. oryzae dia* BGC^7^ are compiled in Fig. 4.

**Figure 4.**
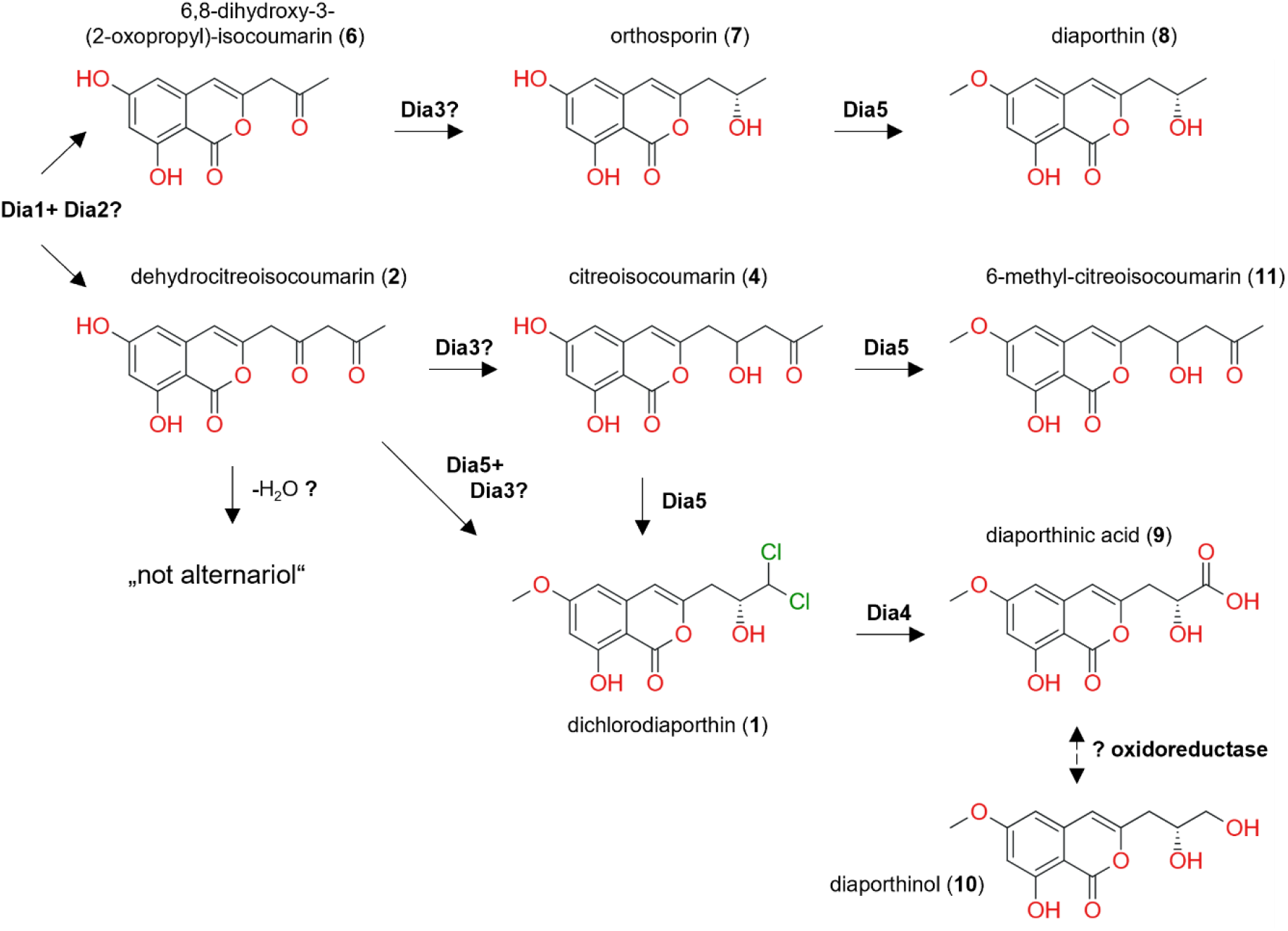
Proposed biosynthetic pathways towards diaporthin (**8**), diaporthinic acid (**9**), and diaporthinol (**10**), all originating from the *dia* BGC in *T. reesei*.

Further, we speculate that the production of **10** might be the product of a reduction of **9**, as **10** arises only downstream of Dia5 (see above) and the relation of **9** and **10** abundance remains similar in all strains (Fig. 3). This reduction might be catalyzed by an unspecific oxidoreductase present in the proteome of *T. reesei*. At this point, this is just speculation; the possible reaction is therefore included in Fig. 4 with dashed arrows.

In this study, we could attribute further metabolites to the *T. reesei dia* BGC *i*.*e*., compounds with the same exact masses as **5, 7** and **8**. Comparison to a commercial standard revealed that the presumed detected compound **5** is actually not **5**, despite exhibiting the same exact mass. This “not alternariol” seems to co-occur together with **2** and to some extent also **7** (Fig. 3). All three compounds accumulate in the absence of Dia4 or Dia5 in comparison to the OEdiaR1 strain, suggesting that there is backlog in the biosynthetic pathway. “Not alternariol” shows a mass difference of 18.01 Da compared to **2**, which suggests that “not alternariol” might be a dehydration (−H^2^O) derivate of **2**.

7 and **8** were previously reported to be the product of the *aoi* BGC in *A. oryzae*^6,7^. In this study, we could attribute compounds with the matching masses to the *dia* BGC (Fig. 3). In the Δdia5 strain, we observed an accumulation of **6** and **7**, and a complete abolishment of **8** production, matching the results from Liu et al.^7^. There, AoiQ was shown to be responsible for the methylation of **7** yielding **8**. We speculate that *T. reesei* Dia1 is able to release both, **2** and **6**, in contrast to *A. oryzae* DiaA, which was reported to only release **2**^7^. Interestingly, in the absence of Dia2, the signals for the assumed **6** and **7** were lower compared to the OEdiaR1 strain, and **8** was not detectable anymore. Thus, T. *reesei* Dia2 might support Dia1 preferably during the release of **6**. The suggested pathway is represented in Fig. 4.

However, we detected all metabolites except **8** in the Δdia2 strain. As the release of **2** and **6** is highly unlikely to occur spontaneously, we must assume that another enoyl reductase complements Dia2. A BLAST analysis^19^ against the predicted *T. reesei* proteome using the *A. oryzae* DiaB protein sequence as query on the JGI genome portal returns an alternative for Dia2 (protein ID 60671^12^, Fig. S5A), a protein containing an uncharacterized subgroup of MBL-fold metallo hydrolase domain (cd07722) and a beta-lactamase associated winged helix domain (pfam17778).

Analogously, the SDR Dia3 was not essential for the biosynthesis of **1** and **9**, which is in stark contrast to the observations by Liu *et al*.^7^. The deletion of *dia3* in the OEdiaR1 strain did not halt the biosynthetic pathway, but the relative abundances of some intermediates changed. Especially the supposed products of Dia3, **6** and **11**, are relatively more abundant compared to the OEdiaR1 strain. For DiaC, a BLAST analysis^19^ as described above using the *A. oryzae* DiaC as query returned several hits (Fig. S5B); the top 5 are all SDRs, like DiaC and Dia3. One or more of them might also accept **2** as substrate and catalyze the necessary reduction reaction in the absence of Dia3.

Although the role of Dia3 in the biosynthetic pathway could be compensated successfully, we observed a strong reduction of the biomass (Fig. S4) and an upregulation of all genes encoding for enzymes in the *dia* BGC compared to the OEdiaR1 strain (Fig. 2, Additional File 2). This might be an indication of a defective negative feedback loop in the absence of Dia3 or due to the gene replacement of *dia3* with the hygromycin resistance marker. Dia3 might also be necessary for self-resistance.

However, Dia3 (protein ID 123964^12^) was previously described to be associated with cellulase signal transduction (Genebank CF653652)^20^. Therein, *dia3* transcripts were detected in elevated amounts shortly after a sophorose pulse, and during cultivation on glucose, and to some extent also on glycerol, but not on cellulose directly^20^. This expression suggests that Dia3 is not associated with cellulose degradation, but it might be involved in other processes. Taken together with the fact that we could detect all metabolites of *dia* BGC in the *dia3* deletion strain further suggests that Dia3 and the other SDRs might fulfill also different biological roles in *T. reesei* and are likely to compensate for each other.

**8** was first isolated in 1953 and was identified to contribute to chestnut blight, to cause wilting of tomato shoots, and to have antibacterial activity against *B. subtilis*^21^. In 1988, **7** was isolated and found to cause necrosis in corn, crabgrass, and soybean^22^. In 2018, both compounds were tested against several bacteria and yeast and found to be bioactive against *Staphylococcus aureus* (even MRSA), *Micrococcus luteus, E*.*coli*, and *S. cerevisiae*^23^. Importantly, the authors describe that the isolated compounds were less active than a crude extract and suggested that **7** and **8** act synergistically with each other and with other compounds.

The bioactivity of **1** was assessed in previous studies and found to be not toxic against zebrafish embryos^24^ or the MDA-MB-435, HepG2, HCT116, H460, and MCF10A cancer cell lines^25^ but it “showed antibacterial activities against *S. aureus, B. subtilis, E. coli, Klebsiella pneumoniae*, and *Acinetobacter calcoaceticus* with MIC values between 25 and 50 μg/ml”^25,26^. Further, it exhibited weak antifungal activity against *Colletotrichum musae* and *Rhizoctonia solani*^27^. In this study, we tested **9** against *E. coli, B. subutilis, S. cerevisisae*, and *A. nidulans*, but could not detect antibacterial or antifungal activity (Fig. S3). To our knowledge there are no reports on the bioactivity of **10**. Given that **8** and **7** act phytotoxic and are suggested to act synergistically^22^, we are prompted to speculate that **9** and **10** might also contribute to this synergistic bioactivity (may it be against plants or microorganisms). However, this is mere speculation at this point and needs to be tested in the future.

## Materials and Methods

### Fungal strains and cultivation conditions

All *T. reesei* strains (Table 3) were maintained on malt extract (MEX) agar plates (30 g/l malt extract, 1 g/l peptone, 15 g/l agar) at 30 °C. Uridine and hygromycin were added if necessary to the final concentrations of 5 mM and 75 µg/ml, respectively. The strains were cultivated in Mandels-Andreotti Medium^28^ (KH_2_PO_4_ 2 g, (NH_4_)_2_SO_4_ 1.4 g, Urea 0.3 g, FeSO_4_·7H_2_O 0.005 g, MnSO_4_·H_2_O 0.0016 g, ZnSO_4_·7H_2_O 0.0014 g, CoCl_2_ 0.002 g, MgSO_4_·7H_2_O 0.3 g, CaCl_2_ 0.3 g, peptone 0.75 g) with 10 g/l glycerol as carbon source at 30 °C at 180 rpm in an orbital shaker. Samples were taken after 48 hours for RNA extraction and biomass determination or 72 hours for metabolite extraction and analyses. The mycelium for RNA extraction was stored at -80 °C. For biomass determination, the mycelium was dried at 80 °C over night until completely dry and weight on a precision scale. The culture supernatants for metabolite analyses were stored at –20 °C.

**Table 3.**
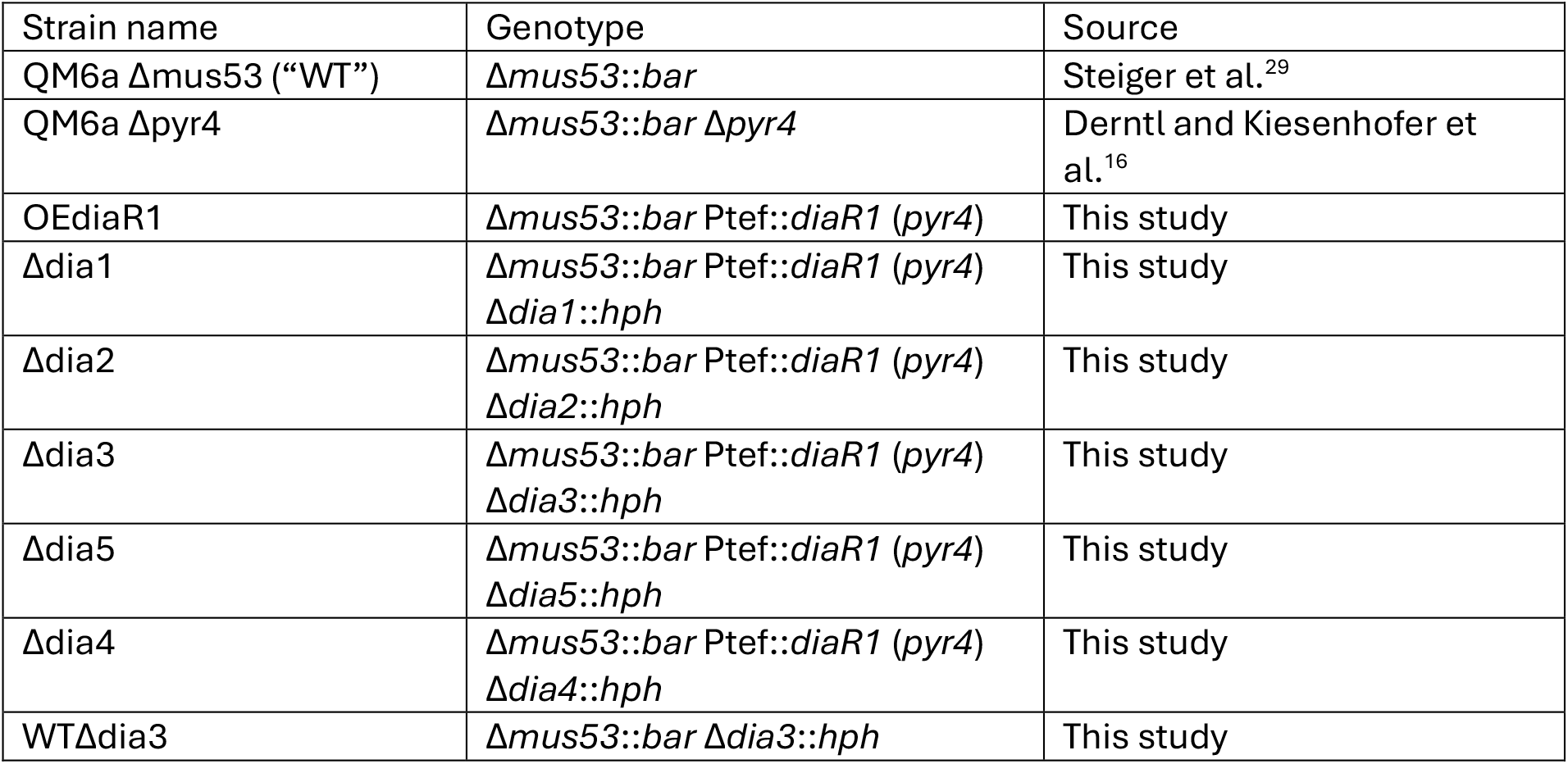
*T. reesei* strains used in this study.

### Fungal transformation

*T. reesei* was transformed using a polyethylene glycol (PEG)-mediated transformation of protoplasts. Spores of the recipient strain were plated on cellophane sheets on MEX plates and incubated overnight at 30 °C. Mycelium was scraped, transferred into 15 mL Buffer A (1.2 M sorbitol, 100 mM KH_2_PO_4_, pH 5.6) with 600 mg Vinotaste Pro (Novozymes, Bagsværd, Denmark) and 0.5 mg chitinase (Sigma-Aldrich), and incubated at 60 rpm and 30 °C for 2–3 hours until protoplasts were released. The suspension was filtered (70 µm sieve), chilled on ice, filled up to 40 mL with ice-cold 1.2 M sorbitol and centrifuged at 2,500 g at 4 °C for 10 min. Then, washed in 30 ml ice-cold 1.2 M sorbitol. The protoplasts were resuspended in 1 mL Buffer B (1 M sorbitol, 25 mM CaCl2, 10 mM Tris-HCl, pH 7.5).

For transformation, DNA (20 µg linearized plasmid or 5 µg fusion PCR products) was mixed with 100 µL protoplast suspension and 100 µL 20 % PEG (mixture of 6.7 mL Buffer B and 3.3 mL “60 % PEG” (60 g PEG 4000, 1 mL 1 M Tris-HCl pH 7.5, 1 mL 1 M CaCl2, 38 mL ddH2O)). After 30 minutes on ice, 60 % PEG was added incrementally, followed by incubation at room temperature for 20 minutes. Buffer C (1 M sorbitol, 10 mM Tris-HCl, pH 7.5) was added gradually, and the mixture was diluted to 50 mL with warm selection medium (containing 1 M sucrose) before plating. Plates were incubated at 30 °C under light for up to a week, and colonies were purified by spore streaking on selection plates containing 0.1 % (v/v) Igepal CA-630.

### Strain construction strategies

The sequences of all used primers are given in the supplemental material. For the construction of *T. reesei* OEDiaR1, the coding region of *diaR1* (protein ID 111742) was amplified with the primers 111742_fwd-AflII and 111742_rev-SpeI and cloned into the plasmid pRP4-TX^30^, putting it under the control of the constitutive *tef1* promoter. The plasmid was linearized by digestion with NotI and transformed into *T. reesei* QM6a Δpyr4, selecting for uridine prototrophy on Mandels-Andreotti Medium^28^ without peptone and 1 % (w/v) glucose. In the resulting strain, OEdiaR1, the *pyr4* locus is reestablished and the DiaR1 expression cassette inserted upstream of the *pyr4* promoter (Fig. S6).

For the deletion of *dia1-5*, a split marker strategy was used. In brief, the 5’ and the 3’ flanks of the genes were fused to overlapping fragments of the hygromycin marker, *hph* from pAN7-1^31^, using a splicing by overlap extension (SOE) PCR (Fig. S7). The two fusion PCR products were simultaneously transformed into the recipient strain (QM6a Δmus53 or OEdiaR1). The hygromycin resistance gene, *hph*, is assembled by a crossing-over between the overlapping regions (Fig. S7), resulting in hygromycin resistance. The gene deletions were confirmed by suitable PCR assays (Fig. S8-S13).

### Genotyping

Mycelium was harvested and pressed dry between two sheets of filter paper. Approximately 50 mg were lysed in 1 mL CTAB buffer (1.4 M NaCl, 100 mM Tris-HCl pH 8.0, 10 mM EDTA, 2 % CTAB, 1 % polyvinylpyrrolidone) with 0.37 g small glass beads, 0.25 g medium glass beads and one large glass bead in a 2 mL screw cap reaction tube using a Fast-Prep-24 (MP Biomedicals, Santa Ana, CA, USA) at 6 m s-1 for 30 sec. The samples were incubated at 65 °C for 20 min and finally centrifuged at 12,000 g for 10 min. The supernatant was transferred to a 2 mL reaction tube and the DNA was purified by a phenol-chloroform-isoamyl alcohol extraction, followed by a chloroform extraction. The samples were then treated with RNase A (Thermo Fisher Scientific) according to the manufacturer’s instructions and the DNA finally precipitated using isopropanol, washed in 1ml 70 % (v/v) ethanol, and dissolved in 10 mM Tris-HCl pH 8.0 after drying.

All PCR reactions for genotyping were performed with the OneTaq DNA Polymerase or the Q5 DNA polymerase (both NEB) according to the manufacturer’s instructions.

### Transcript analyses

Mycelium was harvested and pressed dry between two sheets of filter paper, frozen in liquid nitrogen, and stored at -80 °C for up to a week. Approx. 50 mg of mycelium were disrupted in 1 mL RNAzol RT (Sigma) with 0.37 g small glass beads, 0.25 g medium glass beads and one large glass bead in a 2 mL screw cap reaction tube using a Fast-Prep-24 (MP Biomedicals) at 6 m s-1 for 30 sec. The samples were centrifuged at 12,000 g for 10 min, the supernatant was transferred to a 1.5 mL reaction tube and mixed with ethanol 1:1. The RNA was purified using the Direct-zol RNA MiniPrep Kit (Zymo Research, Irvine, CA, USA) according to the manufacturer’s instructions.

Notably, this kit contains a DNase treatment step. The total RNA was reverse transcribed using the LunaScript RT SuperMix Kit (NEB) according to the manufacturer’s instructions. The cDNA was diluted 1:50 in ddH2O and 2 µL were used as template in a 15 µL reaction using the Luna Universal qPCR Master Mix (NEB) on a Rotor-Gene Q (Qiagen, Venlo, Netherlands). Primers were added and PCR reaction conditions were chosen according to the manufacturer’s instructions. To calculate the relative transcript abundance, we used the Pfaffl method^32^ and the *sar1* gene as reference gene^33^. We also measured the transcript levels of *act1* but they fluctuated strongly (Additional File 2); *act1* was therefore excluded as reference gene.

### Compound isolation and NMR

The supernatant obtained from 300 ml culture was lyophilized and the obtained residue was taken up again in 50 mL water. The obtained clear, aqueous solution was extracted several times with ethyl acetate, until no further extraction of UV-active compounds was observable via TLC (in total 300mL). The collected organic phases were combined, dried over Na_2_SO_4_, filtered, evaporated under reduced pressure and the obtained crude residue was analyzed to proof successful extraction using a Shimadzu Nexera UHPLC system, equipped with an SPD-M20A PDA-detector and a LCMS-2020 single quadrupole mass spectrometer. Chromatographic separation was performed on a Waters XSelect CSH C18 XP column, with an inner diameter of 3.0mm, 50mm length and 130 Å pore size. Buffer system A (water + 0.1 % formic acid) and B (pure acetonitrile) were used at a flow rate of 1.7 mL/min, starting with 5 % B for 0.15, followed by an increase from 5 to 98 % B over a period of 2.05 min and holding at these conditions for 0.30 min.

Preparative separation was performed on a BüchiPure C850 Flashprep, equipped with a DAD, using a Waters SunFire Prep C18 OBD column, with an inner diameter of 30 mm, 100 mm length and 100 Å pore size. Chromatographic separation used buffer A (water + 0.1 % formic acid) and buffer B (acetonitrile + 0.1 % formic acid) employing the following solvent program at a flow rate of 20 mL/min: After injection, 25 % B was applied for 2 min, then it was raised to 40 % B over a period of 7 min, followed by an isocratic separation at 4 0% B for another 7 min. Then, over 30 min the composition was raised to 95 % B and again held at these conditions for 5 min. Fractions were collected and analyzed via the Shimdazu Nexera UHPLC system to check for the purity of the fractions.

Respective fractions containing diaporthinic acid were evaporated under reduced pressure, dried on high vacuum and taken for NMR measurements at 297 K on a Avance III HD 600 spectrometer and processed with standard software. Obtained spectra were calibrated to the residual solvent peak^34^.

### MIC assays

The antibacterial and antifungal MIC assays were performed in 200 µl reactions in a 96-well plate with flat bottom. First, a serial dilution of diaporthinic acid was prepared in DMSO with the following concentrations: 12.8, 6.4, 3.2, 1.6, 0.8, 0.4, 0.2, 0.1 mg/ml. These solutions were added to double strength RPMI 2 % G medium (RPMI 1640 with L-glutamine without sodium bicarbonate (Thermo Scientific), 20.8 g/l; MOPS, 69.06 g/l; glucose, 36 g/l; pH adjusted to 7.0 with NaOH) for the antifungal assays or Müller-Hinton broth for the bacterial assays. 100 µl of these spiked media were transferred to induvial wells and then 100 µl of the respective bacterial or fungal inoculums were then added, in quadruplicates.

The *E. coli* FDA strain Seattle 1946 (ATCC 25922) and *B. subtilis* strain Marburg (type strain, DSM 10) inoculums were prepared by cultivating the bacteria on LB plates (10 g/l peptone, 5 g/l yeast extract, 10 g/l NaCl) at 37 °C over night. Several distinct colonies were suspended in Müller-Hinton broth and the cell density adjusted to an OD600 of 0.02. *The S. cerevisiae* inoculum was prepared by incubating the strain Sa-07140 (type strain, DSM 70449) on YPD agar (20 g/l peptone, 20 g/l glucose, 10 g/l yeast extract) at 30 °C for 48 hours. Several distinct colonies were suspended in sterile, distilled water. The *A. nidulans* inoculum was prepared by cultivating *A. nidulans* FGSC A4 on potato dextrose agar plates at 30 °C for 1 week. Spores were harvested using sterile cotton swabs and resuspended in sterile, distilled water. *S. cerevisiae* and *A. nidulans*, the cell or spore density was adjusted to an OD530 of 0.1. The bacterial MIC assays were incubated at 37 °C for 24 hours, and the antifungal assays at 30 °C for 24 hours. Finally, the optical density was measured on a Promega GloMax microplate reader (Promega) at 600 nm (*E. coli* and *B. subtilis*) or at 560 nm (*S. cerevisiae* and *A. nidulans*).

### HPLC-MS/MS analysis

LC-MS grade acetonitrile (ACN) and LC-MS grade formic acid were acquired from VWR chemicals (Radnor, PA, USA). Water (H2O) was purified in-house using a Barnstead™ Smart2Pure™ Water Purification System from Thermo Fisher Scientific (Waltham, MA, USA). Alternariol (CAS # 641-38-3) standard, Catalog # C5061 (Batch No. 1), was purchased from APExBIO (Houston, TX, USA).

The frozen culture media were thawed, vortexed, and 2 mL of each strain and quadruplicate were transferred into fresh Eppendorf tubes. These tubes were spun down for 2 minutes at 20,000 g at room temperature. From the supernatant, 10 µL were taken out and diluted 1:10 in 2 % ACN + 0.1 % FA. The injection volume was 1 µL.

For analysis of the NMR-confirmed diaporthinic acid standard, a stock solution of the compound in DMSO was diluted 1:10 in 2 % ACN + 0.1 % FA. The commercial alternariol standard (1 mg) was dissolved in 1 mL of 50% ACN (with two drops of NH_4_OH) and further diluted 1:100 in 2 % ACN + 0.1 % FA (c = 10 ng µL-1). The injection volume for both standards was 1 µL.

After dilution of the culture media and the reference standards, the samples were measured employing an untargeted metabolomics workflow in positive ionization mode on a Bruker timsTOF Pro equipped with a VIP-HESI source (Bruker Corporation, Billerica, MA, USA). The frontend was a Thermo Fisher Scientific Vanquish H UHPLC with a Waters Acquity BEH C18 column (150 mm × 1 mm ID, 1.7 μm; Waters Corporation, Milford, MA, USA). Mobile phase A was 0.1 % formic acid in water and solvent B acetonitrile containing 0.1 % formic acid. The following gradient was employed at 40 °C and a constant flow rate of 100 µL min-1: 0 min, 2 % B; 9 min, 98 %; 12 min, 98 % B; 12 min, 2 % B, followed by 4 min re-equilibration. The timsTOF Pro mass spectrometer was operated without trapped ion mobility spectrometry (TIMS off). For positive ionization mode, source capillary voltage was set to 4500 V and dry gas flow to 8 L min-1 at 230 °C. Sheath Gas Flow was set to 4.0 L min-1 at a temperature of 200 °C, with active exhaust being activated. Scan mode was set to Auto MS/MS with 12 Hz MS spectra rate and 16 Hz MS/MS spectra rate, resulting in a total cycle time of 0.5 s. Scan range was set from 20 to 800 *m/z*.

Individual compounds were quantified on MS1 level employing the open-source application Skyline version 23.1^35,36^ normalizing to the TIC, based on exact mass only.

## Supporting information

BGC genebank files

RT-qPCR data

MIC assay

## Author’s contribution

IB performed the HPLC-MS/MS analysis and co-drafted this manuscript.

SL performed the compound isolation and identification and co-drafted this manuscript.

KP, LF, PA, and LTSK constructed different fungal strains, performed RT-qPCR analyses and cultivations.

RF, CS were involved in the compound isolation and identification.

FR, RBG, RLM, ARMA provided resources and were involved in supervision.

MS was involved in the study design and supervision.

CZ was involved in the study design and supervision, co-drafted this manuscript and performed the MIC assays.

