## Supplemental Figures 1-13 for "*In vivo* activation of the *dia* BGC allows consolidation of the biosynthetic pathways of diaporthin, dichlorodiaporthin, diaporthinic acid, and diaporthinol"

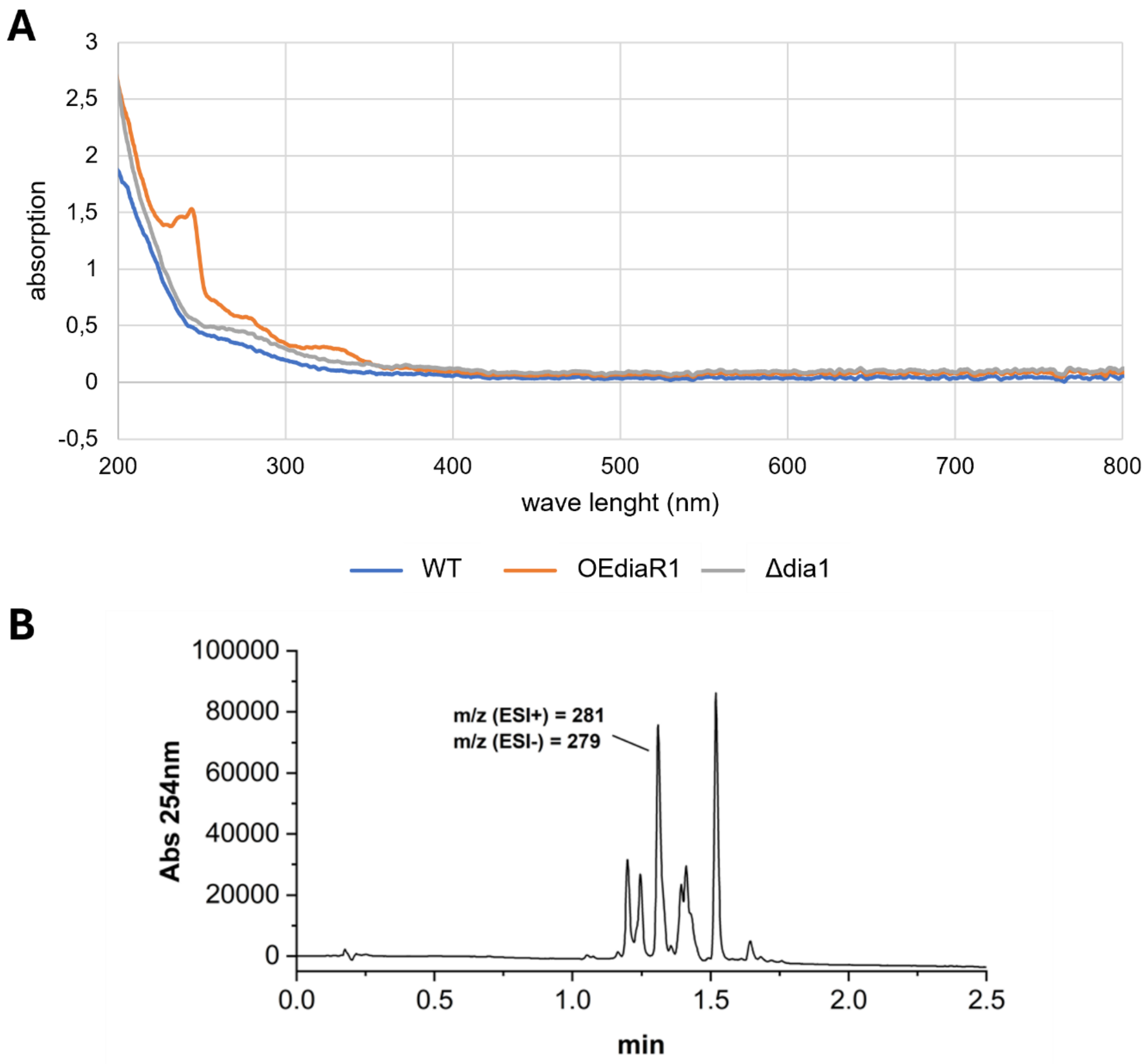


**Figure S1.** (**A**) The T. reesei strains QM6a Δmus53 (WT), OEdiaR1, and Δdia1 were cultivated in MAM + glycerol for 72 hours and the absorption spectrum of the resulting supernatant measured. (**B**) The supernatant of OEdiaR1 was subjected to a HPLC-PDA/MS analysis.

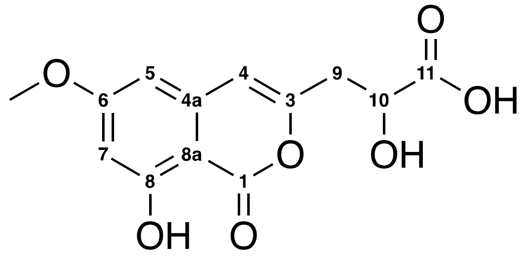


**Figure S2.** 1D- and 2D-NMR of the peak with *m/z* (ESI+) = 281 and *m/z* (ESI-) = 279 in Fig. S1B.

1H-NMR (600MHz, d6-DMSO): δ = 2.73 (dd, J = 8.6, 14.7 Hz, 1H, H9-A), 2.90 (dd, J = 4.4, 14.7 Hz, 1H, H9-B), 3.85 (s, 3H, OCH3), 4.30 (dd, J = 4.4, 8.6 Hz, 1H, H10), 6.53 (d, J = 2.3 Hz, 1H, H7), 6.57 (s, 1H, H4), 6.61 (d, J = 2.3 Hz, 1H, H5) ppm

13C-NMR (600MHz, d6-DMSO): δ = 37.86 (C9), 55.96 (OCH3), 67.60 (C10), 99.39 (C8a), 100.52 (C7), 101.30 (C5), 106.05 (C4), 139.39 (C4a), 154.11 (C3), 162.52 (C8), 165.29 (C1), 166.52 (C6), 174.40 (C11) ppm

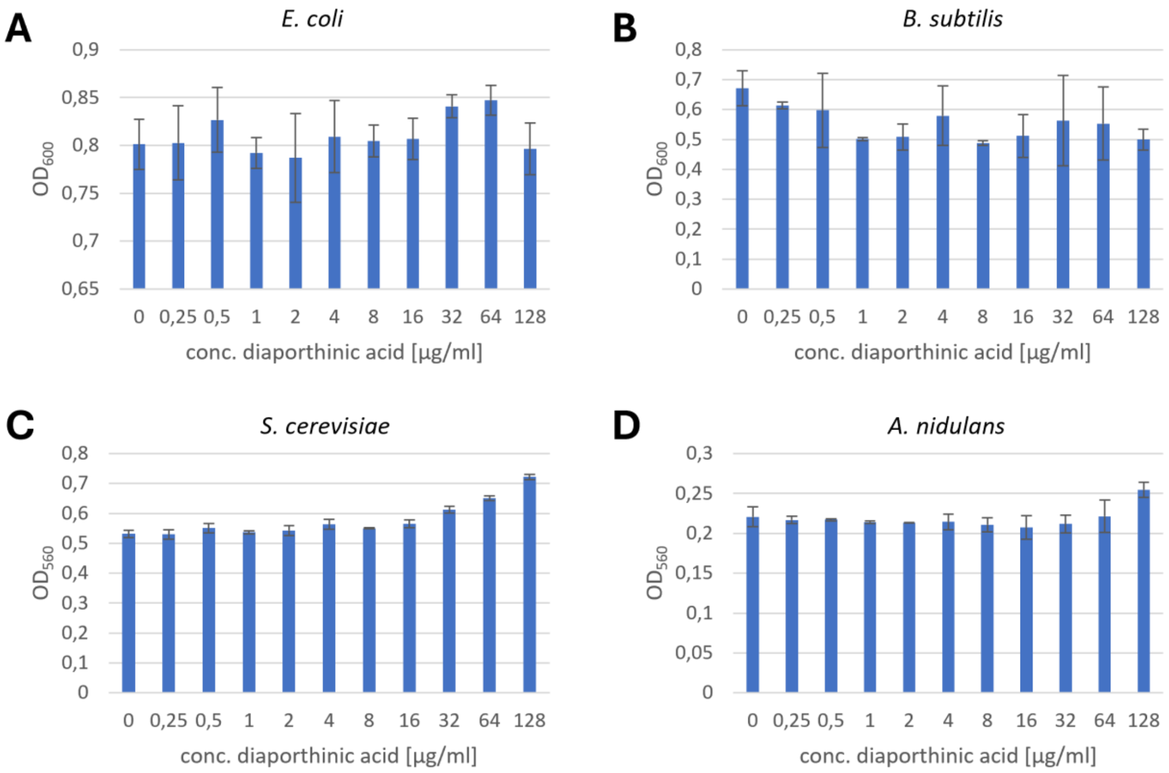


**Figure S3.** The indicated microorganisms were tested from sensitivity against diaporthinic acid in a MIC assay. The obtained culture density is shown in dependency of the diaporthinic acid concentration.

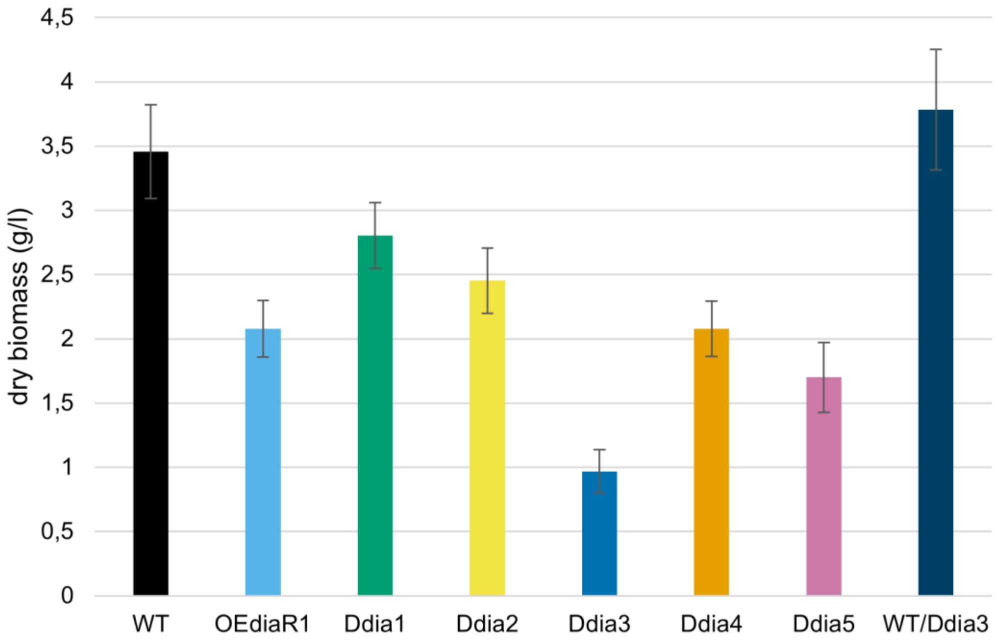


**Figure S4.** The indicated *T. reesei* strains were cultivated in MAM+glycerol for 48 hours and the resulting biomass harvested and dried. The measurement was performed in quadruplicates, the error bars indicate the standard deviation.

**A**

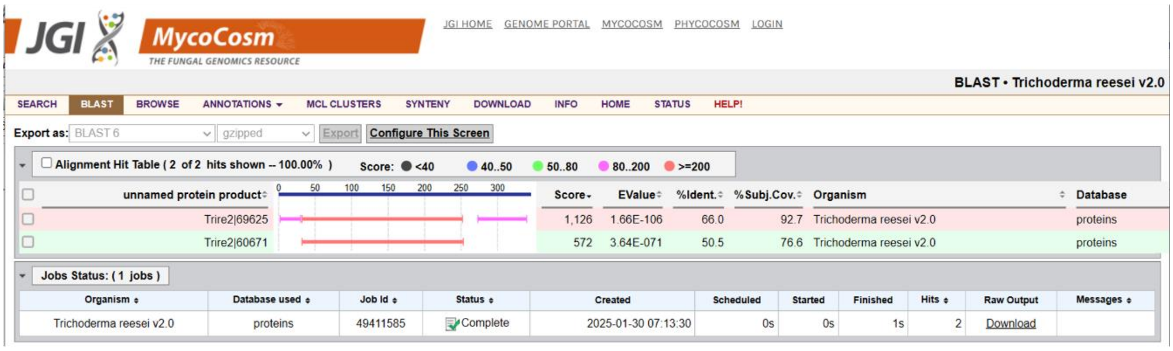


**B**

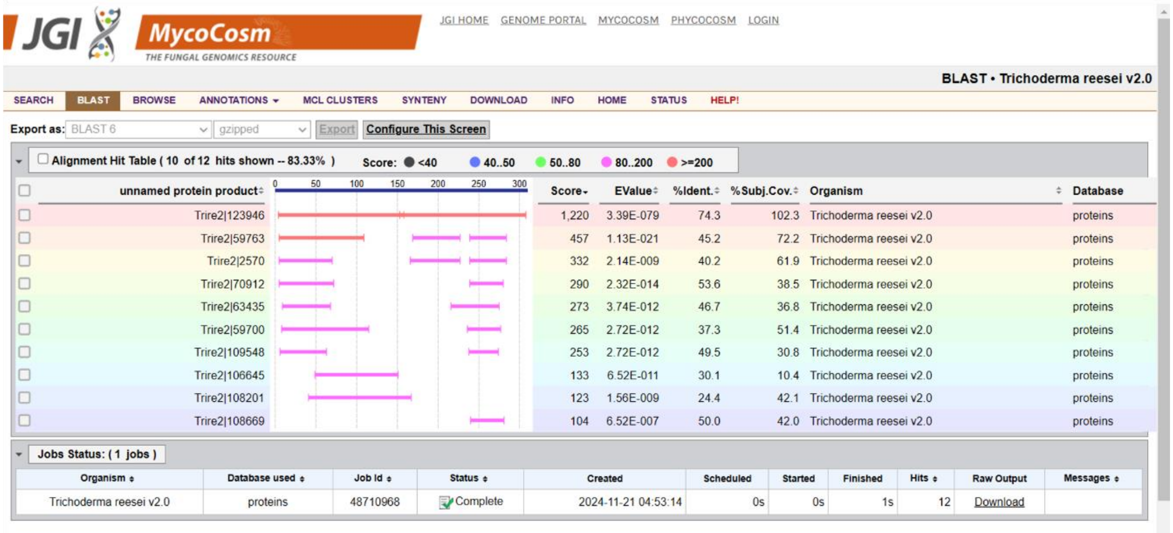


**Figure S5.** The protein sequences of *A. oryzae* DiaB (**A**) and DiaC (**B**) were used as query in a BLAST analysis against the proteome of *T. reesei* on https://mycocosm.jgi.doe.gov/Trire2/Trire2.home.html

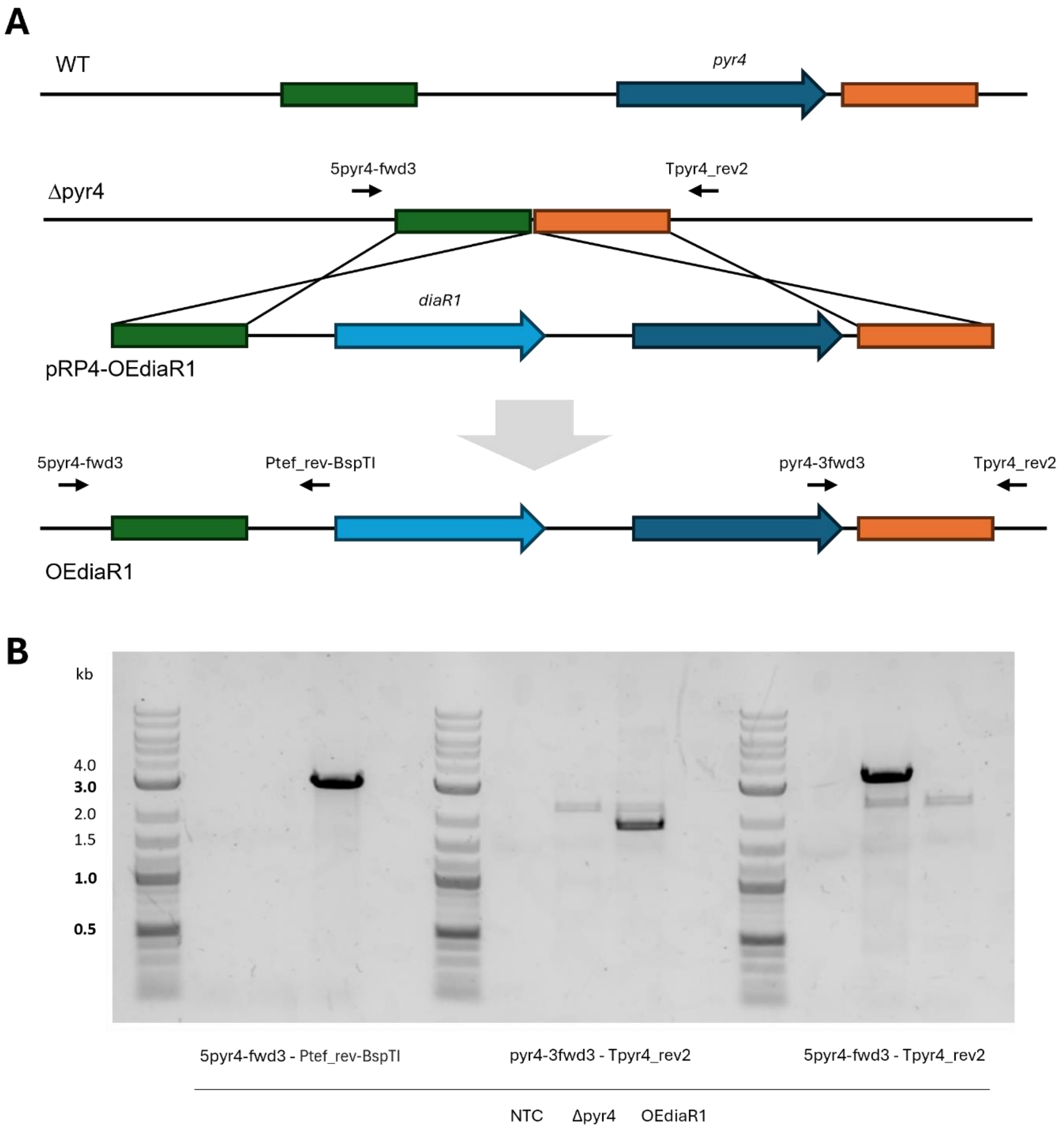


**Figure S6. Construction of *T. reesei* OEdiaR1. (A)** The plasmid pRP4-OEdiaR1 was linearized and inserted into *T. reesei* Δpyr4. Following a double cross-over, the expression cassette was inserted at the *pyr4* locus while simultaneously re-establishing the *pyr4* locus. The black arrows indicate the position of the primers used for genotyping. **(B)** The chromosomal DNA of the indicated strains was isolated and used as template in PCR assays using the indicated primers. NTC, no template control.

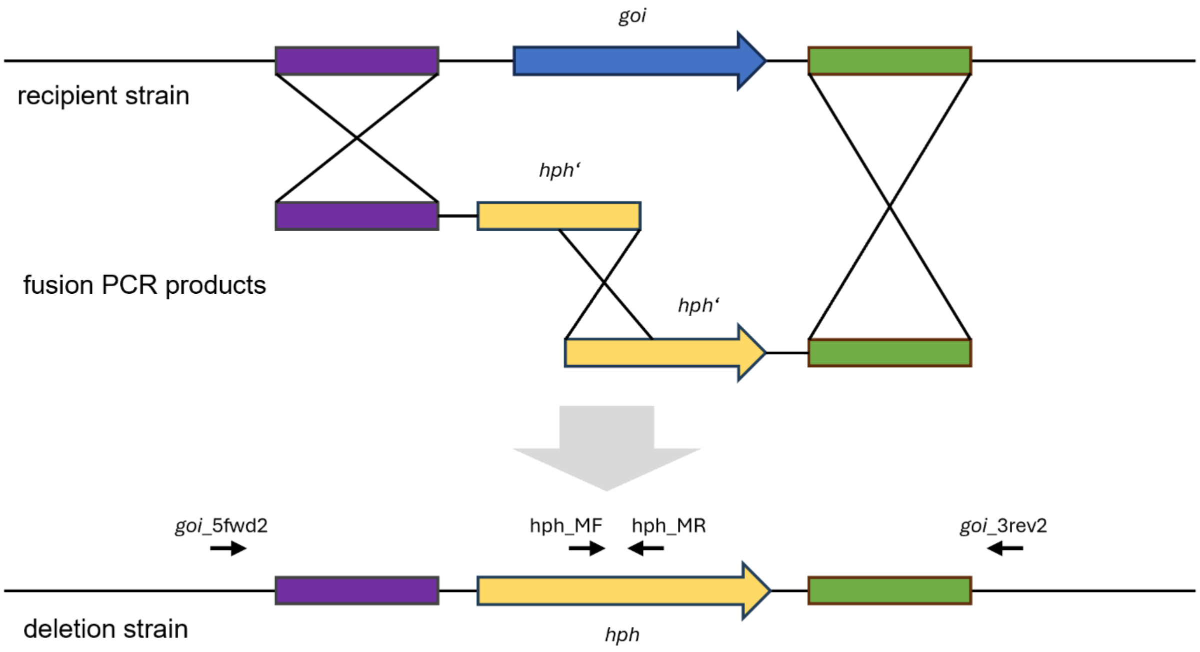


**Figure S7. Split marker strategy for gene deletions.** For the deletion of gene of interest (goi), two distinct fusion PCR products were constructed *in vitro* using a splicing by overlap (SOE)-PCR. The first PCR product contains the 5'flank and the first two thirds of the maker gene *hph*, while the second PCR product consists of the last 2 thirds of *hph* and the 3'flank. Upon a triple crossover, the marker is assembled and integrated at the correct locus, resulting in the deletion of the *goi* and its replacement with the *hph* marker. The black arrows indicate the positions of the primers used for genotyping in Fig. S8-S13.

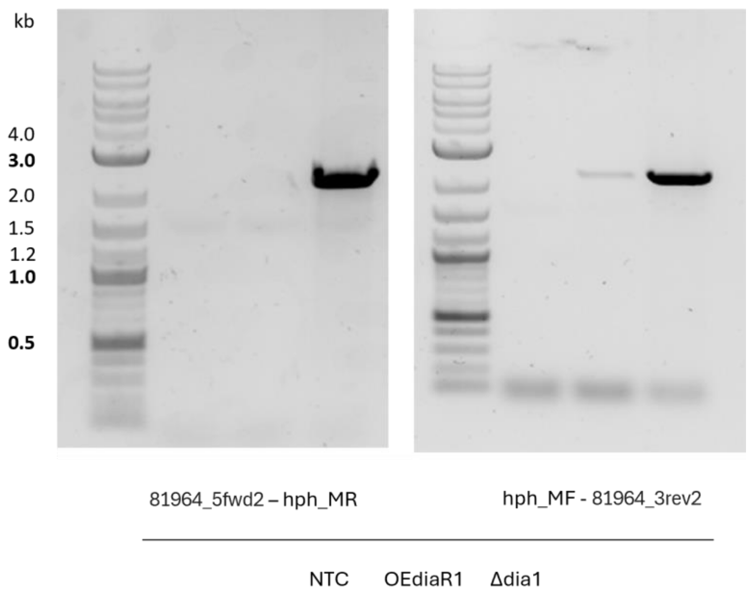


**Figure S8. Genotyping of *T. reesei* Δdia1.** To verify the replacement of *dia1* with the *hph* resistance cassette as depicted in Fig. S7, the chromosomal DNA of the indicated strains was isolated and used as template in PCR assays using the indicated primers. NTC, no template control.

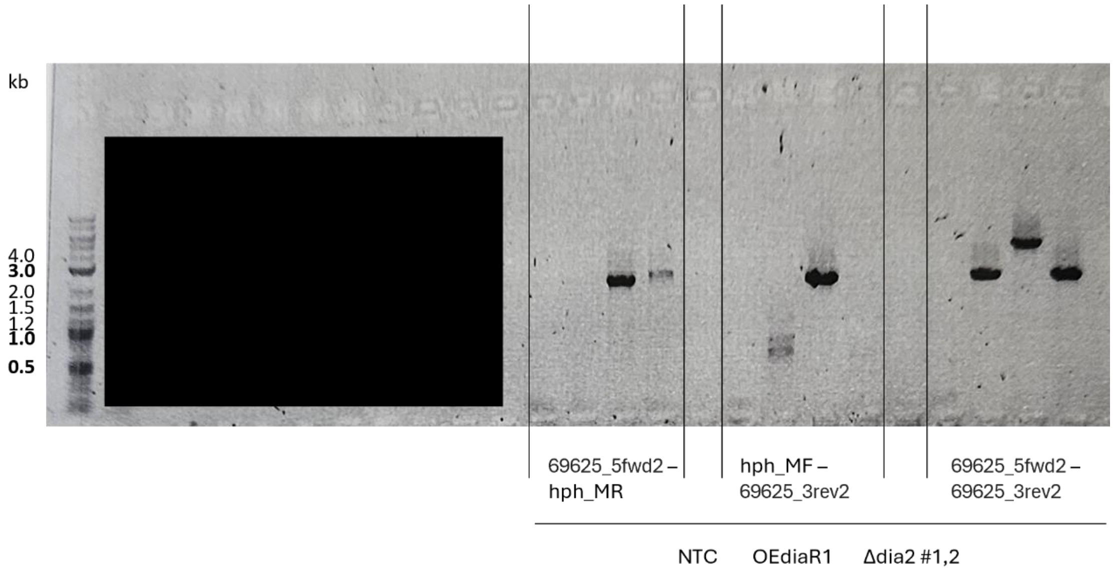


**Figure S9. Genotyping of *T. reesei* Δdia2.** To verify the replacement of *dia2* with the *hph* resistance cassette as depicted in Fig. S7, the chromosomal DNA of the indicated strains was isolated and used as template in PCR assays using the indicated primers. NTC, no template control.

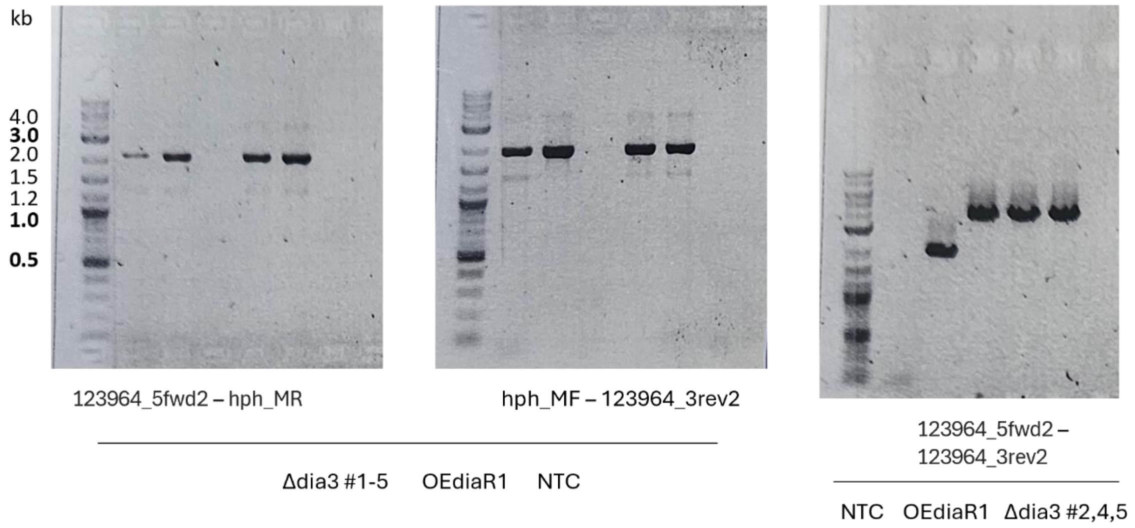


**Figure S10. Genotyping of *T. reesei* Δdia3.** To verify the replacement of *dia3* with the *hph* resistance cassette as depicted in Fig. S7, the chromosomal DNA of the indicated strains was isolated and used as template in PCR assays using the indicated primers. NTC, no template control.

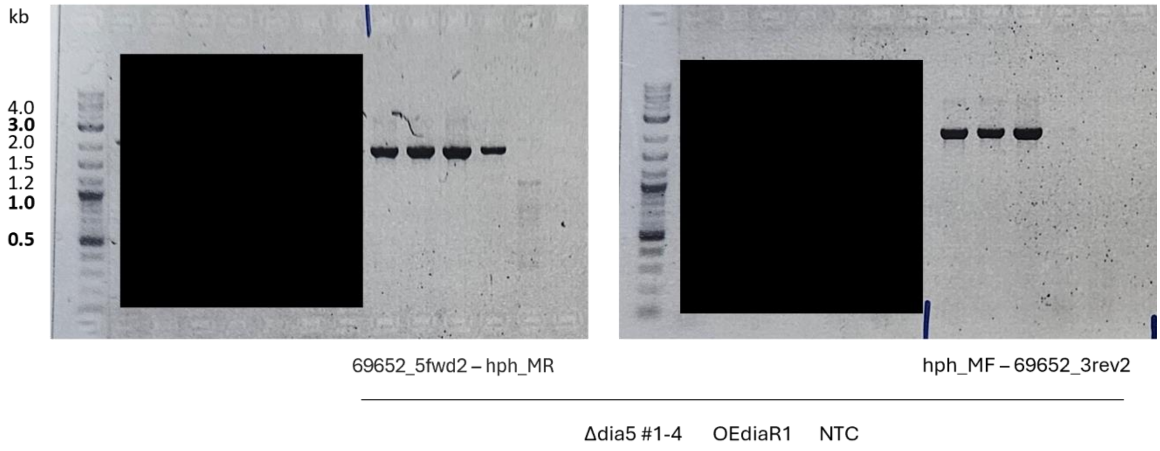


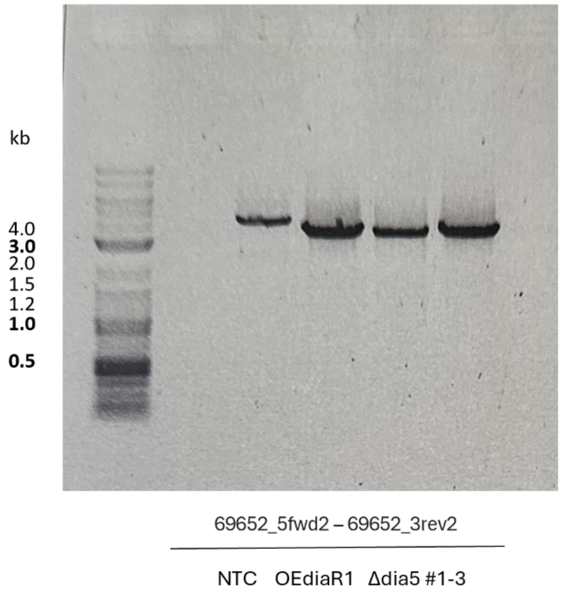


**Figure S11. Genotyping of *T. reesei* Δdia5.** To verify the replacement of *dia5* with the *hph* resistance cassette as depicted in Fig. S7, the chromosomal DNA of the indicated strains was isolated and used as template in PCR assays using the indicated primers. NTC, no template control.

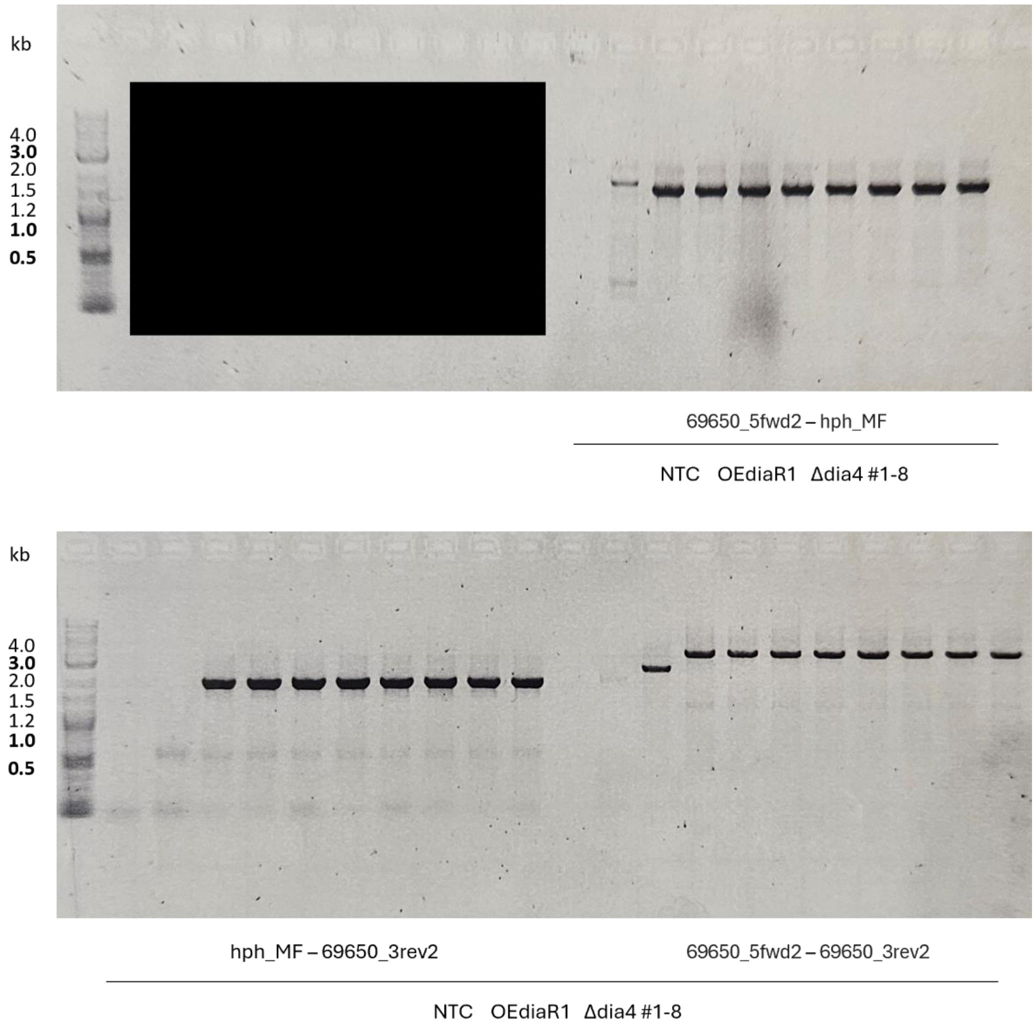


**Figure S12. Genotyping of *T. reesei* Δdia4.** To verify the replacement of *dia4* with the *hph* resistance cassette as depicted in Fig. S7, the chromosomal DNA of the indicated strains was isolated and used as template in PCR assays using the indicated primers. NTC, no template control.

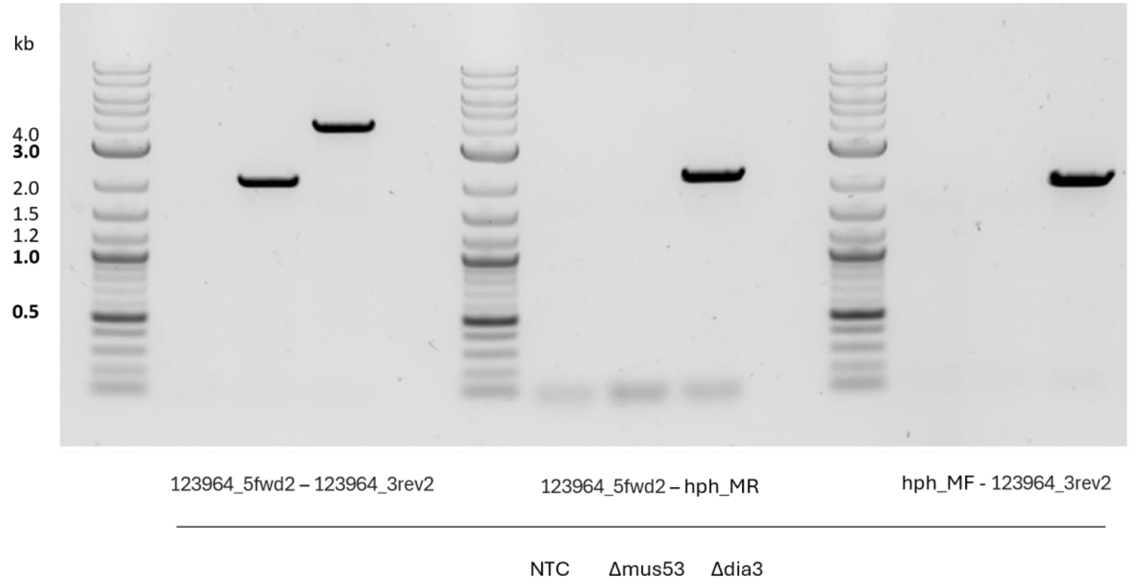


**Figure S13. Genotyping of *T. reesei* WT/Δdia3.** To verify the replacement of *dia3* with the *hph* resistance cassette as depicted in Fig. S7, the chromosomal DNA of the indicated strains was isolated and used as template in PCR assays using the indicated primers. NTC, no template control.

**Table S1.** Homologs of the *T. reesei dia* BGC genes in *D. pomorum*

| Gene name | Protein ID | Enzyme class | Homolog in *D. pomorum* |
|---|---|---|---|
| *dia1* | 81964 | polyketide synthase | KAL1641334 |
| *dia2* | 69625 | beta-lactamase-like | JAKJXN020000121 (22,623 – 23,120) |
| *dia3* | 123964 | dehydrogenase | KAL1641336 |
| *dia5* | 69652 | bifunctional flavin-dependent halogenase / methyltransferase | KAL1641335 |
| *dia4* | 69650 | the FAD-dependent oxioreductase | KAL1641337 |
| *diaR1* | 111742 | zinc cluster protein | JAKJXN020000121 (15,791 - 14,892 |
